# Isolated correlates of somatosensory perception in the mouse posterior cortex

**DOI:** 10.1101/2022.07.16.498499

**Authors:** Michael Sokoletsky, David Ungarish, Yonatan Katz, Ilan Lampl

**Affiliations:** Department of Brain Sciences, Weizmann Institute of Science, Rehovot, Israel

## Abstract

To uncover the neural correlates of stimulus perception, experimenters commonly use tasks in which subjects are repeatedly presented with a weak stimulus and instructed to report, via movement, if they perceived the stimulus. The difference in neural activity between reported stimulus (hit) and unreported stimulus (miss) trials is then seen as potentially perception-related. However, recent studies found that activity related to the report spreads throughout the brain, calling into question to what extent such tasks conflate perception-related activity with report-related activity. To isolate perception-related activity, we developed a paradigm in which the same mice were trained on both a regular go/no-go whisker stimulus detection task and a reversed contingencies version, in which they reported the absence of a whisker stimulus. By comparing no-report trials across the two tasks, we located perception-related activity within a posterior network of cortical regions contralateral to the stimulus. In addition, we found this activity was on average an order of magnitude lower than report-related activity and began after the low-level stimulus response. In summary, our study revealed the mouse cortical areas associated with the perception of a sensory stimulus independently of a perceptual report.

## Introduction

How sensory stimuli become perceived is one of the greatest mysteries in neuroscience. At its most fundamental level is the question of why is it that some stimuli are perceived at all, enabling the subject to report on their presence, while other stimuli remain subliminal. To answer this question, the classical approach, known as the go/no-go detection task, has been to repeatedly deliver a weak stimulus, which is sometimes perceived and sometimes not, and instruct the subject to report if the stimulus was perceived. The report consists of an action that can range anywhere from a verbal indication in humans (Del Cul et al., 2007) to pressing a button in non-human primates (de Lafuente et al., 2006) and licking a water sprout in mice (Fig. 1A; Sachidhanandam et al., 2013). Neural activity is then compared between reported and unreported stimulus trials. From the resulting difference, these studies have identified several brain regions as possible substrates of perception, including the primary sensory cortex (Sachidhanandam et al., 2013; Takahashi et al., 2016), secondary sensory cortex (Y ang et al. 2016; Kwon et al., 2016), premotor cortex (Lafuente and Romo, 2005; Lafuente and Romo, 2006), and prefrontal cortex (Van Vugt et al., 2018).

**Figure 1:**
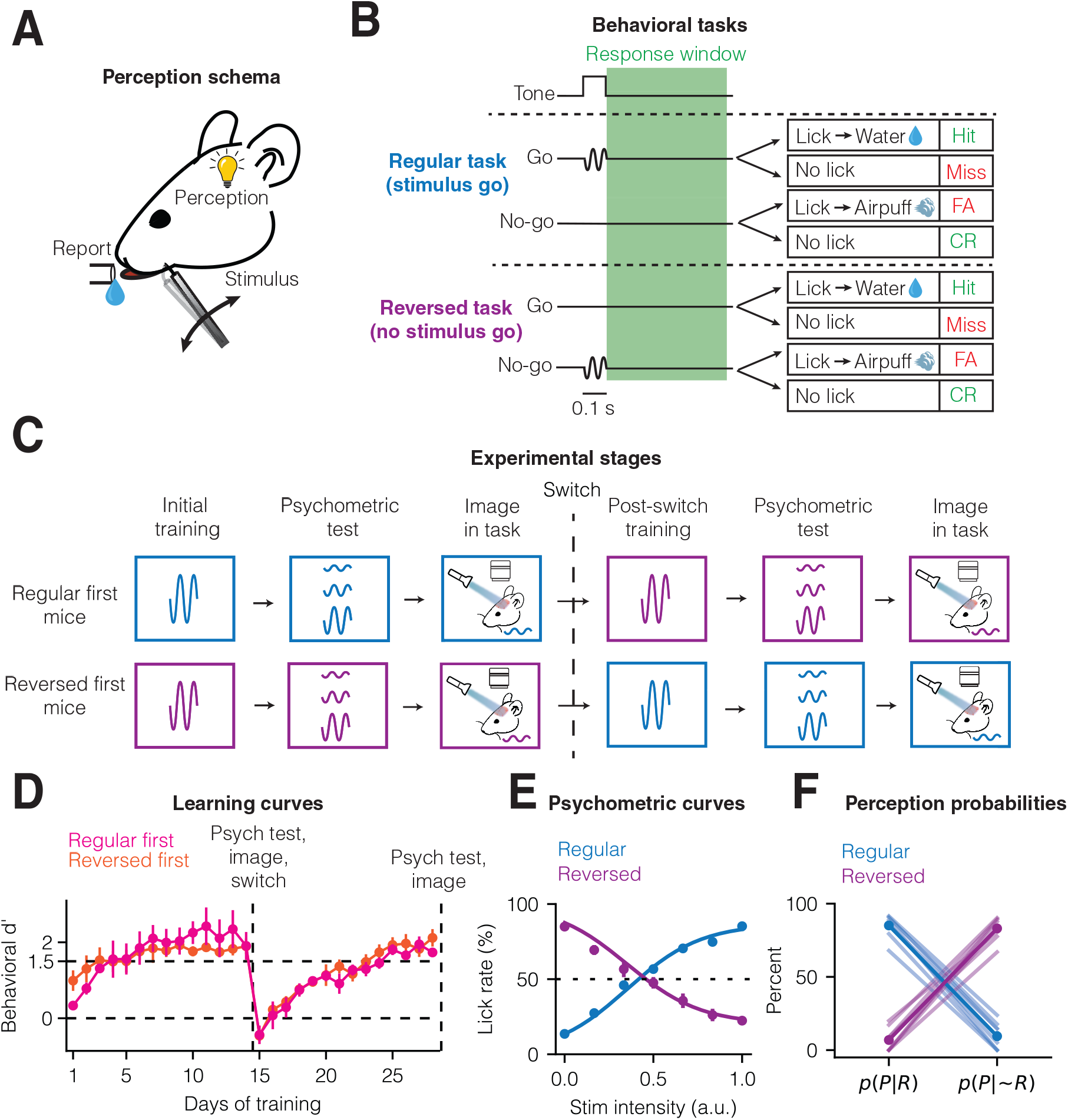
Mice similarly learn and perform the regular go/no-go task and its reversed version. **(A)** Schematic of the experimental set-up. **(B)** Trial structure and outcome types across the two go/no-go tasks. **(C)** Training structure, with mice belonging to either regular first (n = 4) group or the reversed first (n = 4) group. Each group had analogous training other than the initial and reversed contingencies. **(D)** Learning curves across the two tasks for each of the two groups. **(E)** Psychometric curves for each task, with the lick rate as a function of stimulus intensity (normalized to 1). **(F)** Results of Bayesian analysis of the psychometric curves for the probability of perception given the mice made or did not make a report in trials within each task. Mean ± s.e.m in plots.

Recently, however, studies have shown that activity related to the report spreads throughout the brain, even to sensory regions (Orsolic et al., 2021; Steinmetz et al., 2019; Zatka-Haas et al., 2021; Musall et al., 2019). Comparing activity between trials with and without a report may thus yield not only the neural correlates of perception, but also the neural correlates of the report itself, with no clear way of separating the two (Zagha et al., 2022). Compounding this issue is the fact the latter can include not only activity driving the report’s execution, but also preparation for the report, sensory feedback during the report, and, if there is a reward or punishment in the task, anticipation of the report’s consequence. In addition, it can involve activity related to uninstructed movements that tend to co-occur with the report, e.g. trunk and limb movements (Musall et al., 2019). We will refer to all these factors collectively as *report-related activity*.

Several recent studies in mice were able to separate the stimulus response from report-related activity within stimulus discrimination tasks by modeling the effect of each variable on neural activity (Steinmetz et al., 2019; Musall et al., 2019). These studies found that while report-related activity is dominant across the cortex, the stimulus response still explains a considerable portion of the activity in its primary sensory region. However, because in discrimination tasks, unlike in detection tasks, mice compare stimuli that are perceived to some extent, the stimulus response presumably consists of a mixture of perception-related activity as well as activity that results from the stimuli regardless of perception, which we will refer to as the *low-level stimulus response*. It thus remains unclear to what extent cortical activity in response to a stimulus is perception-related, separately from both the low-level stimulus response and report-related activity.

To isolate perception-related activity, we trained the same mice on two go/no-go detection tasks: one corresponding to the standard design and a reversed contingencies version in which the mice were to lick if they did not perceive the stimulus. This allowed us to obtain pairs of trials across the tasks in which the stimulus was identical, but whereas in the regular task the report indicated the mice perceived the stimulus, in the reversed task, the report indicated the mice did not perceive the stimulus. Hence, this allowed us to obtain the correlates of perception independently from both stimulus and report by comparing no-report trials across the tasks. We then leveraged widefield calcium imaging to look for perception-related activity throughout the cortex. Using both grand average and decoding analyses, we discovered that isolated perception-related activity appears within a posterior network of cortical regions contralateral to the stimulus and characterized its spatiotemporal dynamics in relation to the low-level stimulus response and report-related activity.

## Results

We trained mice in one of two go/no-go tasks which we termed regular and reversed (Fig. 1A, n = 8 mice total, n = 4 regular task then reversed task and n = 4 reversed task then regular task). In the regular task, the mice learned to make a report by licking a water sprout when presented with a 20 Hz deflection of the left C2 whisker and a simultaneous auditory tone (both lasting 100 ms) and not to make a report when the tone was given without the deflection (Fig. 1A,B). Correct reports were rewarded with sweetened water, and incorrect reports were punished with a light airpuff. In the reversed task, the mice learned to make a report when the tone was given alone, and to not make a report when it was concurrent with the whisker deflection. A retractable lickport was presented to the mouse at the start of the response window and retracted at the end of every trial to avoid confounds from early licking. The response window began at the end of the tone and lasted 500 ms. Importantly, reward (in hit trials) or punishment (in false alarm trials) was given at the end of the response window and regardless of when the mice licked, so that the neural activity within the response window would not be affected by the reward or punishment.

To train the mice, we used a strong whisker deflection for two weeks of daily sessions, at which point we proceeded only with the mice that had reached expert level on the initial task (discriminability index *d’* > 1.5, Fig. 1C,D, Supplementary Fig. 1A). We then constructed a psychometric function for whisker stimulation using a range of intensities and fitted this psychometric function with a sigmoid to determine the perceptual threshold, i.e. the stimulus intensity at which the mice responded correctly 50% of the time (Fig. 1E). After recording neural activity during the task with the stimulus intensity adjusted to this threshold, we switched the tasks for each group of mice and repeated these steps, taking care that every task parameter other than the contingency remained the same (see Methods). During initial training, mice in each group began to consistently perform at expert level after about five training days. After switching, the mice ultimately reached similar performance to the initial task, albeit at a slower rate of about 10 days. We also returned a subset of mice (n = 2 regular first and n = 2 reversed first) to the initial task, which took longer than the initial learning but a similar duration to the post-switch learning (Supplementary Fig. 1B).

From the psychometric curves, we can calculate the theoretical probability that a mouse perceived the threshold stimulus given that it made a report, *p*(perceivedļreported), and the probability that it perceived the threshold stimulus given that it did not make a report, *p*(perceivedļ~reported) (Methods). In the regular task, *p*(perceivedļreported) = 85% and *p*(perceived|~reported) = 10%, while in the reversed task *p*(perceived|reported) = 7% and *p*(perceived|~reported) = 83% (Fig. 1F). Thus, we can reasonably interpret the mouse’s report as indicating that it either perceived or did not perceive the threshold stimulus depending on the task. As further evidence that the near-50% lick rate for threshold stimuli was due to trial-to-trial variability in the perception of the stimulus rather than random licking, cutting the whiskers in trained mice abolished correct responses to threshold stimuli (n = 2 regular task and n = 2 reversed task, Supplementary Fig. 1C).

Comparing the lick latencies across the tasks, we found that they were considerably lower for reversed hits and false alarms (mean latencies 110 ± 11 ms and 112 ± 10 ms, respectively; Supplementary Fig. 1D) than for regular hits and false alarms (mean latencies 147 ± 28 ms and 154 ± 29 ms, respectively). This suggests that it was faster for mice to make the determination that there was no stimulus, even when it was erroneous in the case of reversed false alarms.

We recorded neural activity during each task by performing widefield calcium imaging throughout the cortex. The mice were transgenic, expressing GCaMP6s in excitatory neurons in all layers. Fluorescence was measured through the intact, cleared skull, and then aligned to the Allen Mouse Common Coordinate Framework (CCF v3; Fig. 2A,B, Supplementary Fig. 2). We first compared activity between regular hits and regular misses (Fig. 2C, “classical comparison”). In both trial types, focal activity was clearly visible in the C2 barrel in S1 by 100 ms from the start of stimulation. Subsequently, there was increased activity across the cortex, but to a greater degree in hits. This was reflected in a positive difference throughout the cortex between hits and misses (Fig. 2C, bottom row), which was especially pronounced in a set of medial regions consisting of the upper and lower limb and trunk somatosensory cortices (collectively, the limb-trunk cortex) and the primary motor cortex (Fig. 2D).

**Figure 2:**
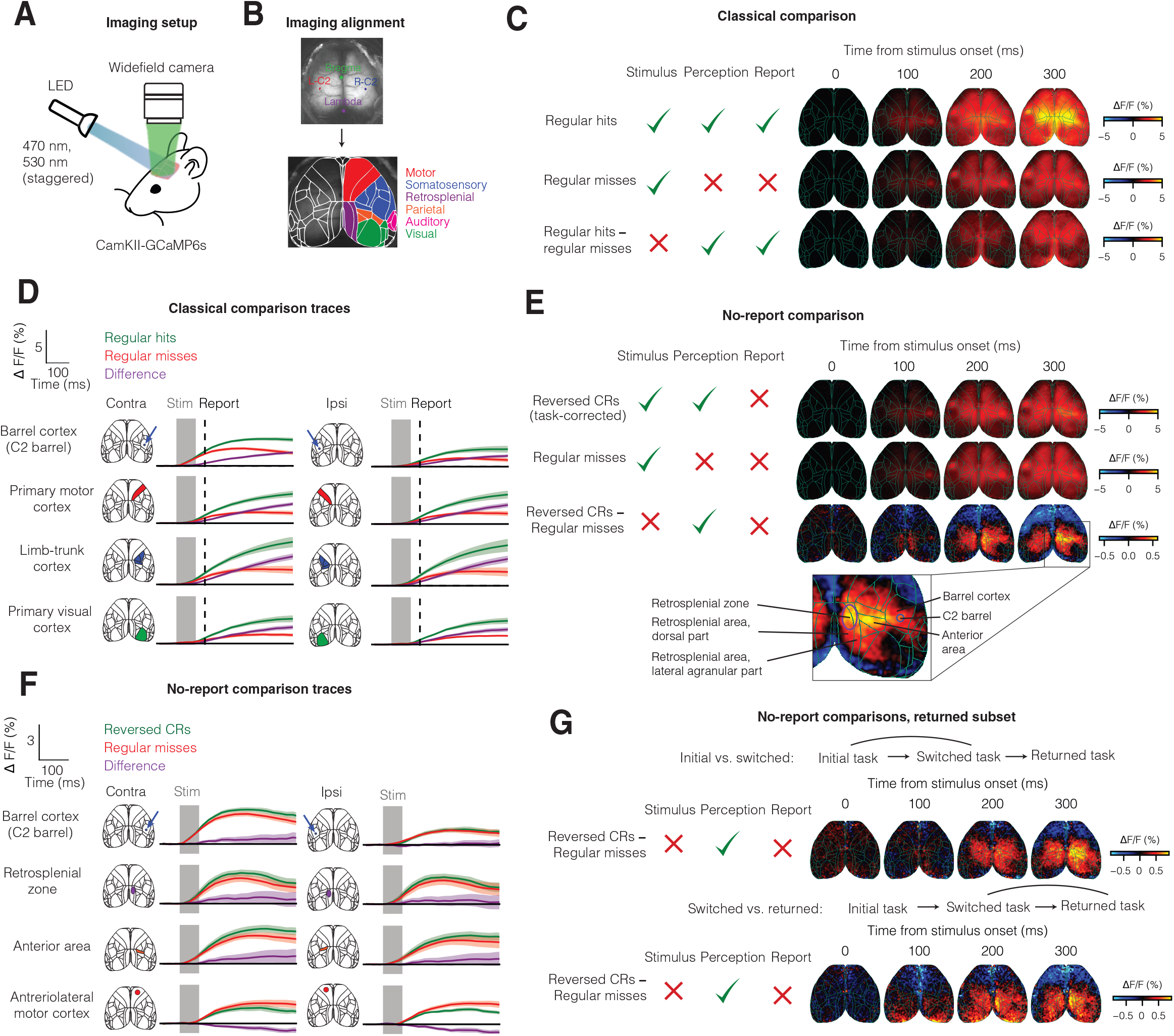
Isolated correlates of perception in the posterior cortex. **(A)** Widefield calcium recording set-up. **(B)** Selection of four points for alignment to the Allen CCF v3: bregma, lambda, and left and right C2 barrels (top) and the results of alignment, labeled by regional association (bottom). **(C)** Grand averaged timelapses of, from top to bottom, activity during regular hits, regular misses, and the difference between them. **(D)** Averaged traces of activity across four key cortical regions for regular hits, regular misses, and the difference between them. **(E)** Grand averaged timelapses of, from top to bottom, activity during reversed correct rejections, regular misses, and the task-corrected difference between them. **(F)** Averaged traces of activity across four key cortical regions for regular CRs (task-corrected), regular misses, and the difference between them. **(G)** Control for time/experience-dependent changes in activity. (Top) Task-corrected no-report comparison between the switched task and initial task (n = 4 mice), (bottom) task-corrected no-report comparison between the returned task and the switched task (n = 4 mice). Mean ± s.e.m in plots.

To interpret the differential maps, we assume that, in general, differences in activity across the trial types in the two tasks consist of three factors: the low-level sensory response, perception-related activity, and report-related activity. Since regular hits and regular misses differ by both report and perception, it is unclear which of the two is responsible for the observed difference in neural activity. To isolate perception, we would instead like to compare reversed correct rejections to regular misses. In both trial types the mice received a whisker stimulus and did not make a report, but in reversed correct rejections they did not make a report because they perceived the stimulus, whereas in regular misses, they did not make a report because they did not perceive the stimulus. Thus, the difference between the two should be limited to perception alone.

When comparing trial types across the tasks, however, we observed an unexpected phenomenon: a marked difference between neural responses across tone-alone trials from each task (regular correct rejections and reversed misses), which was similar across other trial type pairs in each mouse (Supplementary Fig. 3A). This difference, which we termed the “task factor”, was highly correlated between sessions pairs within the same mouse but only weakly across mice (Supplementary Fig. 3B), reflecting diversity in its appearance: in some mice posterior regions were more active in the reversed task, while in others they were less active, and in some the differences where symmetrical across the hemispheres while in others they were asymmetrical. Further analysis showed that the task factor was neither correlated to the much smaller time or experiencedependent activity changes in the returned mouse subset (Supplementary Fig. 3C), nor was it affected by controlling for uninstructed movements of the body and whisking (Supplementary Fig. 3D-F; Methods).

Assuming that the task factor is indeed common to all trial types, the simplest way to correct for it is to subtract it from any difference between the tasks. Only upon doing so did we discover a difference between reversed correct rejections and regular misses which was similar across mice and differed from each mouse’s task factor (Supplementary Fig. 4A, Fig. 2E). This difference, which presumably represents purely perception-related activity, was broad: it spread throughout the posterior cortical regions of both hemispheres, but was stronger in the hemisphere contralateral to the stimulus (Fig. 2E,F). The location of its peak was on average within the retrosplenial cortex and anterior area (Fig. 2E, bottom) with some variance across mice (Supplementary Fig. 4A).

Could it be that this difference is not a true correlate of perception, but instead stems from another difference between the trial types we have not accounted for? We considered three possibilities. First, it could be that it stems from a time or experience-related difference in the brain response across the two tasks, as we previous examined for the task factor. Whereas the above mean map of perception (Fig. 2E) was obtained from all mice, here we tested this possibility in the returned mouse subset alone by comparing the perception correlates calculated from the initial and reversed tasks to those from the reversed and returned tasks (Fig. 2G). Despite the opposite direction of time and experience between these comparisons, we observed similar correlates of perception in each case. Second, even in the absence of a report, differences in uninstructed movement amplitudes between reversed correct rejections and regular misses may account for the observed activity difference. To test this, we selected subsets of trials from each trial type to minimize the difference between the distributions of whisking and body motion across the trial types (Supplementary Fig. 4B,C; Methods) and repeated the comparison, which yielded similar perception correlates (Supplementary Fig. 4D). Lastly, the perception correlates could stem from a perception-independent difference between the task contexts we have not accounted for with the task factor. To test this, we repeated the experience with another group of 8 naïve mice (n = 4 regular first, n = 4 reversed first) that have not undergone training with the strong stimulus and licked randomly (Supplementary Fig. 4E,F). Though these mice exhibited a similar difference between regular hits and regular misses to the trained mice (Supplementary Fig. 4G), they did not exhibit the observed differences between reversed correct rejections and regular misses (Supplementary Fig. 4H).

The above findings concern comparisons of grand averages across trial types, but they may be missing more subtle differences between the response distributions. To examine such differences, we applied a pixel-based binary decoder (Steinmetz et al., 2019; Zatka-Haas et al., 2021; Methods), in which the distribution of single-trials responses within each pixel was compared across trial types in which all factors other than the one examined were constant. This also offered us a statistically powerful way to test the significances of any results as we could compare them to the results obtained when performing the same procedure on data with shuffled trial labels (Methods).

For perception, similarly to before, we compared across the distributions of regular misses and reversed CRs, then subtracted the task component obtained from comparing the distributions of reversed misses and regular CRs. We found the resulting perception-related activity (Fig. 3A,B) was both spatially and temporally highly similar to the one found from the grand average comparison. It first simultaneously emerged in a number of regions in across both hemispheres (primary somatosensory cortex, retrosplenial cortex, and anterior area) at 133 ms (defined as the first frame at which activity became significantly different from zero, p < 0.05, FDR-corrected permutation test), then spread to other primary and higher-order visual cortices (anterior, anteromedial, and posteriomedeial) and somatosensory cortices (barrel, trunk, and limbs) by 200 ms. The decoder approach could also be used to compare perception-related activity directly to low-level stimulus and report-related activities (Fig. 3A). The low-level stimulus response was most prominent in the contralateral (right) S1-bc (barrel cortex) activity starting already at 33 ms. Simultaneously or a little later, it also became significant in neighboring regions in the same hemisphere – R-S1-n (nose, 33 ms), R-S2 (33 ms), R-S1-tr (trunk, 33 ms), R-S1-un (unassigned, 33 ms), R-M1 (33 ms), and R-M2 (67 ms) – as well as regions in the ipsilateral hemisphere – L-S1-bc (barrel cortex, 33 ms), L-S1-n (nose, 33 ms), L-S2 (67 ms), L-M1 (67 ms), and L-M2 (67 ms) (Fig. 3A,B). For report, significantly-encoding regions were M1, M2 and the somatosensory regions in both hemispheres (trunk, upper and lower limb cortices in particular) starting at 33 ms after stimulus onset, and every cortical region by 100 ms (Fig. 3A,B), consistent with the activity difference we observed between hits and misses within the regular task (Fig. 2C).

**Figure 3:**
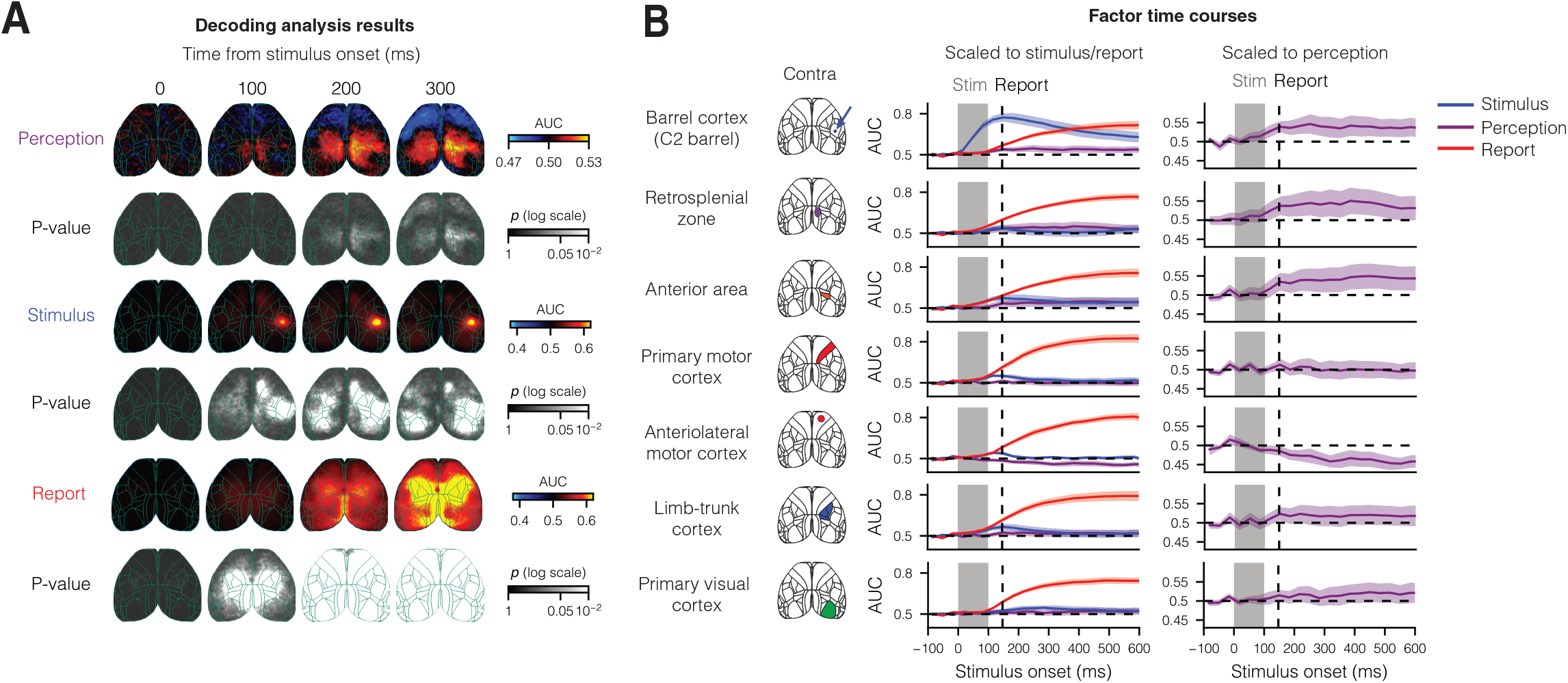
Comparison of perception, stimulus, and report-related activities. **(A)** Decoding AUC maps for stimulus, report, and perception, as well as the P-values for each, over four time bins. Each time bin is labeled by its center. **(B)** (Left) Mean AUCs of perception, stimulus, and report for each of 6 cortical regions in the right hemisphere, scaled to the maximum stimulus or report AUCs. (Right) Mean AUCs of perception, scaled to its maximum, across examined regions: R-S1-C2 (right C2 barrel cortex), R-S2 (right secondary somatosensory cortex), R-M1 (right primary motor cortex), R-M2 (right secondary motor cortex), R-RSPd (right retrosplenial cortex, dorsal part), RSPagl (right retrosplenial cortex, lateral agranular part), and R-PPC (right posterior parietal cortex). Error bands in (B) are s.e.m.

## Discussion

We found that isolated perception-related activity appears in a set of cortical regions that is distinct from the set of regions activated due to the stimulus alone or in a report-correlated manner. While the single whisker stimulus primarily evoked a response in its corresponding barrel, and report-related activity spread throughout the cortex, perception-related activity was mostly restricted to a posterior network of regions, particularly parts of the contralateral retrosplenial cortex and anterior area. Crucially, the magnitude of this activity was much lower than either stimulus-related or report-related activity, at least within the paradigm in which the stimulus is at the threshold of perception.

It is important to note the limitations of this study in terms of the recorded widefield signal, which did not consist of firing of individual neurons, but rather the combination of somatic and neuropil activity of groups of neurons in each pixel. While this signal has been shown in a variety of GCaMP6-expressing mouse lines to be correlated with local spiking activity (Allen et al., 2017; Clancy et al., 2019; Makino et al., 2017, Xiao et al., 2017), it is possible that more heterogeneous local activity, perhaps in a small subset of cells, was averaged out in our recordings. Moreover, comparison with cellular-resolution imaging suggested higher correlation with layer 1 than layer 2/3 signals (Allen et al., 2017), so a perception-related signal located deeper in the cortex is even more likely to be missed. Better recording precision can be obtained using either two-photon imaging or high-throughput extracellular probes (Steinmetz et al., 2021), or widefield calcium imaging in transgenic mice with GCaMP expression localized to deeper layers (Musall et al., 2021; Mohan et al., 2022).

While this study was only correlational in nature, it is tempting to speculate about the causal role of the observed perception-related activity in terms of the mouse’s ability to perceive somatosensory stimuli or even stimuli of other modalities. The anterior area in the mouse cortex largely overlaps the posterior parietal cortex (Harvey et al., 2012), which has been casually examined during perceptual decision-making, but its role remains controversial. Some studies have found no effect when silencing this region during tactile (Guo et al., 2014; Gallero-Salas et al., 2021) or auditory (Erlich et al., 2015) tasks, while others found that its silencing impairs performance during auditory (Zhong et al., 2019) or visual (Zhou and Freedman 2019) tasks. It could be that the source of this discrepancy lies in the nature of the stimuli in the tasks: in the former studies, the stimuli were clear and fixed, whereas in the latter studies they were intentionally ambiguous. Thus, it may be that the PPC plays a specific role in perceiving ambiguous stimuli (Zhong et al., 2019). The role of the retrosplenial cortex is even less clear, and to the best of our knowledge has not been tested in tasks with ambiguous stimuli. However, a few recent studies found it was crucial during visual navigation (Franco and Goard 2021; Kira et al., 2022), and simulation of its layer 5 induced a dissociation state in another study (Vesuna et al., 2020). Future work manipulating activity in the retrosplenial cortex and anterior area using a near-threshold stimulus task can shed valuable light on the causal role of these regions in stimulus perception.

## Materials and Methods

### Animal subjects

All surgical and behavioral procedures were approved by the Institutional Animal Care and Use Committee of the Weizmann Institute of Science. Experiments were conducted with male and female mice between 8 and 24 weeks of age. Two transgenic strains purchased from the Jackson Laboratory were used to create the mouse line used for imaging: CaMKII-tTA (JAX #007004) and TRE-GCaMP6s (JAX #024742). All mouse strains were of C57BL/6J background. For widefield imaging we used 4 CaMKII-GCaMP6s mice. No statistical methods were used to predetermine the sample size. All mice undergoing training were housed alone under an inverted 12 hour light/dark regime and trained in darkness during the dark part of their cycle.

### Surgical procedures

All surgeries were performed under 1–2% isoflurane anesthesia. After the mouse was anesthetized, its scalp was shaved and lidocaine ointment (Emla) was applied to its skin. A circumferential incision was then made to expose as much of the skull as possible. For widefield imaging, after clearing the skull’s periosteum, two small marks were made using nail polish at the positions of bregma and lambda for mapping purposes. We then applied a layer of cyanoacrylate (Krazy Glue) to optically clear the bone, and placed a 6 mm radius coverslip (thickness = 150 μm) to make the surface even. This resulted in clearly visible blood vessels after a few minutes. A headbar was then affixed behind the coverslip using more cyanoacrylate. Finally, a black, 2 mm high lightshield was glued in front of the coverslip to attenuate the light from the widefield imaging reaching the mouse’s eyes.

### Behavior

The behavioral set-up based on a custom LabVIEW program controlling and receiving data from both a NI card (PCIe 6351) and two Arduinos (Mega 2560). The Arduinos were used to create a pulse-width modulated output for the servomotor and speaker, as well as to synchronize the LED colors with the frames during widefield imaging (see “Widefield imaging” below). Eight mice were trained in the regular and reversed go/no-go tasks. Trials were automatically initiated with a 1-2 second baseline period (randomized in each trial) prior to the tone. The tone was a 100 ms long beep at 6 kHz and 71 dB at the mouse’s position. In go trials in the regular task, and no-go trials in the reversed task, the tone was concurrent with a galvanometric stimulation of the C2 whisker. This stimulation was done by inserting the C2 whisker into a 23-gauge metal tube attached to the galvanometer. To ensure stimulation reproducibility, we positioned the tube tip 2 mm from the whisker’s base using XYZ-adjustable stage. Finally, we reversibly attached the whisker to the tube using non-cured silicone (Yellow Body Double, Smooth-On).

Upon the end of the tone, a servo-mounted lickport was presented to the mouse. This was done to prevent any confounds from early licking before or during the stimulation. The lickport was mounted on an XYZ-adjustible stage, and positioned for each mouse such that at the end of the servo’s movement, it was 2 mm below the upper lip and 1 mm posterior to the tip of the lower lip. The mouse had a 500 ms response window to lick before the servo rotated the lickport away. A piezo sensor attached at the base of the lickport sensed the licks. If the mouse licked, then depending on whether it did so during a go or no-go trial, it respectively received a reward or a punishment upon the end of the lick window. The reward consisted of a 7 μL drop of 5% sucrose water, while the punishment was a 500 ms airpuff applied to the face. If the mouse was rewarded, then the lickport stayed for an additional 6 seconds to allow the mouse to fully consume the water. If it was punished, then there was an additional 8 second timeout period. Otherwise, the delay to the beginning of the next trial was 2 seconds.

Animals were trained and experimented on for approximately 2 months. One week prior to the beginning of training, they were water restricted and habituated to the set-up by being provided free water on it. Once habituated, all whiskers on the left and right whisker pads were trimmed other than C2. Before training, whether in the regular or inverted contingency, all trials were go trials and the mice were taught to lick after the tone. The no-go trials were introduced starting at the first day which is displayed in the learning curves. In the first 14 days of learning, mice were trained to perceive a relatively strong whisker stimulation (~6° amplitude, 20 Hz, 100 ms). Initially, go/no-go trials were not fully random, but required the mouse to perform correctly either once or twice (randomized to prevent pattern learning) before switching. Only once the mice reached *d’* = 1.5 did we make go/no-go fully random, at which point we observed that doing so had no detrimental effect on performance. After 14 days of training for each mouse, we performed 2-3 psychometric experiments to determine the perceptual threshold (see below). We then recorded from it using this threshold stimulus. Finally, contingencies were reversed and all the steps starting at the first day of training were repeated.

### Psychometrics

During learning, the discrimination index (*d’*) was calculated for each session as *Z*(*p*(hit))-*Z*(*p*(false alarm)), where *Z*(p) is the inverse of the standard normal cumulative distribution function (cdf), evaluated at the probability *p*. Intuitively, it represents the normalized difference between the hit percentage and false alarm percentage, expressed in standard deviations.

After learning each task but before recording, the stimulus amplitude was varied to create a psychometric curve for each mouse and task. The intensity threshold was deduced by fitting perception performance across stimulus intensities with a logistic function (Psignifit):

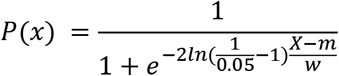

where *x* is stimulus intensity, *P*(*x*) is the report probability, and *m* and *w* are free parameters that were fitted using maximum likelihood estimation (Schütt et al, 2015). Parameter *m* measures the threshold intensity of the psychometric function, and this was the intensity given to the mice on the subsequent recording sessions.

To estimate the probability of perception at threshold stimulation, given that the mouse made or did not make a report, we made two assumptions:

1. The right asymptotic limit of the psychometric curve (at a hypothetical infinite stimulus amplitude) represents the probability that it will make a report if it did perceive the stimulus, *p*(*R*|*P*).
2. Both the above probability and the probability that the mouse will make a report at zero stimulus intensity (i.e., if it did not perceive the stimulus), *p(R|~P)*, are independent of stimulus intensity, i.e., they are the same for the threshold stimulus.

From these probabilities we sought to estimate the probability that the mouse perceived the threshold stimulus if it made a report, *p(R|P)*, and the probability that it perceived the threshold stimulus given that it did not make a report, *p(~R|P)*. To do so, we applied Bayes’ theorem to obtain:

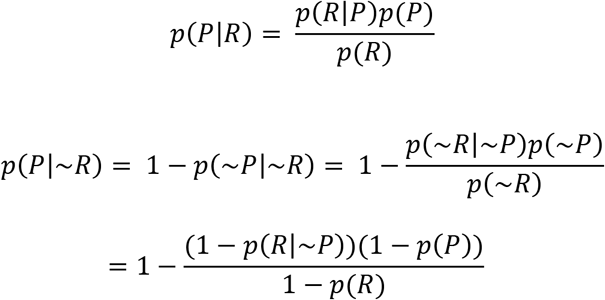

where *p*(*R*) = 0.5 is the probability of licking in response to the threshold stimulus, and *p*(*p*) is the overall probability of perceiving the threshold stimulus. Combining these equations by solving for *p*(*P*) and simplifying yields:

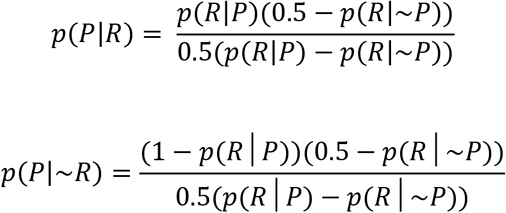

### Video monitoring

Two CMOS cameras (DMK 37BUX287) were used to monitor mouse movements. They were positioned to capture the right side of the mouse’s face and its body. Footage was either recorded at 300 frames per second and downsampled to 30 frames per second, or recorded at 30 frames per second. To illuminate the mouse, we placed one 850 nm LED (M850L3, Thorlabs) in front of the mouse and another 850 nm LED underneath it. Other sources of background illumination (especially the 470 nm and 530 nm light during widefield imaging) were removed by placing 830-865 nm bandpass filters (BP845, MidOpt) in front of the sensor in each camera.

### Widefield imaging

Widefield imaging was done using a macroscopic lens (Nikon AF Micro-NIKKOR 60 mm f/2.8D) mounted on a scientific complementary metal-oxide semiconductor (sCMOS) camera (Quantalux, Thorlabs) running at 60 frames per second. To capture GCaMP fluorescence, a 500 nm long-pass filter (FELH0500, Thorlabs) was placed in front of the camera’s sensor. The total field of view was 11.5 x 9.5 mm^2^ and the image resolution was 230 x 190 after 4x spatial binning (spatial resolution: ~50 μm per pixel). To isolate Ca^2+^-dependent signals, we corrected for intrinsic signals by multiplexing excitation light at two different wavelengths: 470 nm (M470L3, Thorlabs) and 530 nm (M530L3, Thorlabs). This was accomplished by coupling the two LEDs into the excitation path using a 505 nm long-pass dichroic mirror (DMLP505R, Thorlabs), and using the strobe counter output of the camera to alternate illumination between the two LEDs from frame to frame. This resulted in one set of frames with blue and the other with green excitation at 30 fps each. Because excitation of GCaMP with the green LED results in non-calcium-dependent fluorescence, this allowed us to correct each blue frame by dividing by the interpolated green frame:

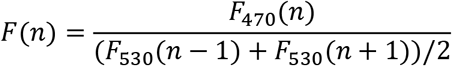

where *n* is the frame number (Ma et al., 2016). This ratio was then normalized to its mean during the baseline period of each trial–the 500 ms immediately preceding tone onset. All widefield data was motion-corrected in the X and Y axes using FFT-based subpixel registration (Guizar-Sicairos et al., 2008). Next, it was rigidly aligned to the Allen CCF v3 (Wang et al., 2020) using the locations of lambda and bregma, marked on the skull during the surgery stage, and the locations of the left and right hemisphere C2 barrels, found by stimulating the left and right C2 whiskers. Alignment was tested by also giving a full-field visual stimulus consisting of 470 nm blue light, and observing that the resulting response overlapped with V1 (data not shown).

### Movement matching

To match uninstructed movements between two different trial types, we manually selected regions of interest (ROIs) for each mouse consisting of the whisking pad (from the face camera) and body including the limbs (from the body camera). From each region we extracted a trace of motion energies (ME) at each frame, defined as the mean of the absolute differences in pixel intensities between the current frame and the previous frame. The traces were averaged within the response window period of each trial type to yield a distribution of whisking and body movements, and the earth mover’s distance (EMD; Yossi et al., 1998) was calculated between each pair of distributions across trial types. We then ran the following iterative algorithm to minimize the overall EMD: (1) calculate the current sum of the whisking and body movement EMDs, (2) for each *i* from 1 to *N*, where *N* is the number of trials in the trial type with the greater number of trials, calculate the hypothetical sum of EMDs if that trial is removed, (3) remove the trial for which the decrease between the current sum and hypothetical sum is the greatest, (4) repeat steps 2-3 until the EMD sum is at an absolute minimum. Note that if the trial types reached parity in the number of trials, the iterations were run successively in one trial type and then the other to maintain parity.

### Decoding analysis

We used a binary decoder to separate decode stimulus, report, and perception (Steinmetz et al., 2019, Zatka-Haas et al., 2021). This decoder was a generalization of classical choice analysis (Britten et al., 1992) and allowed us to combine multiple comparisons across the two tasks. We will describe this method for each of the three factors examined (stimulus, report, and perception).

For decoding the stimulus, we compared regular misses to regular CRs, and reversed FAs to reversed hits. In each of the two comparisons, both report and perception were controlled for: perception was absent in both, and report was absent in the former comparison and present in the latter (with similar latency). We then temporally aligned all trials from each trial type to the stimulus onset, and created separate vectors for the activity of each pixel at each time bin. Our goal was to calculate the area-under-curve (AUC) of the receiver operating characteristic (ROC) for a simple threshold classifier across the two comparisons. Using the relationship between this AUC and the Mann–Whitney *U* for a comparison between two independent groups with *n_1_* and *n_2_* samples:

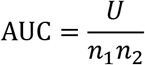

We calculated the AUC at each pixel and time bin as the sum of the *U* statistics for the two comparisons in the numerator divided by the sum of products of the number of trials in each of the trial types in the denominator. In other words, the final AUC was the weighted average of the AUCs from the comparison.

For decoding report, the procedure was the same as above, but with different comparisons: regular FAs vs. regular CRs, reversed hits vs. reversed misses, regular hits vs. reversed CRs, and reversed FAs vs. regular misses. In each one, both stimulus and perception were controlled for across the trial types. For decoding perception, however, a different procedure was used. Rather than performing a weighted average of AUCs, we instead calculated the AUC from the reversed CRs vs. regular misses comparison and corrected it by the AUC from the reversed misses vs. regular CRs comparison (task factor) by direct subtraction.

Significance in each case was measured by comparing the resulting choice probabilities to 0.5 using the Wilcoxon signed-rank test across mice at each pixel and time bin, then FDR-correcting the resulting map of *p*-values at α = 0.05.

---

**Supplementary Figure 1:**
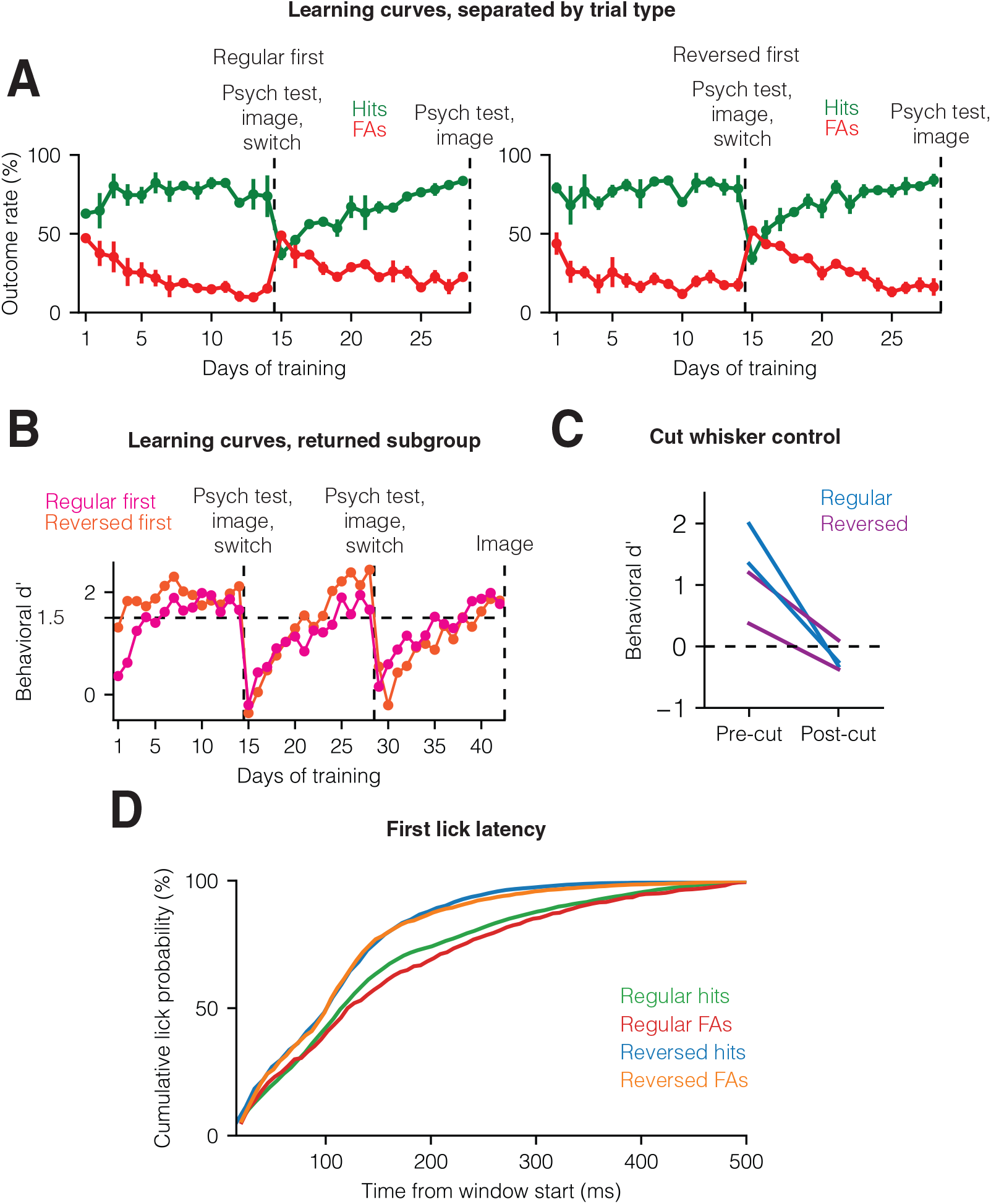
Additional task and group comparisons. **(A)** Mean hits and false alarms across the regular first (n = 4) mice (left) and reversed first (n = 4) mice. **(B)** Mean task performance across the subset of 4 mice (n = 2 regular first, n = 2 reversed first) that we also switched back to the initial task. **(C)** Decrease in behavioral performance to chance level in response to threshold stimulation after cutting the whisker (n = 2 regular first, n = 2 reversed first). **(D)** Cumulative distribution function of the first lick latency across the four trial types, pooled across mice. Mean ± s.e.m in (A) and (B).

**Supplementary Figure 2:**
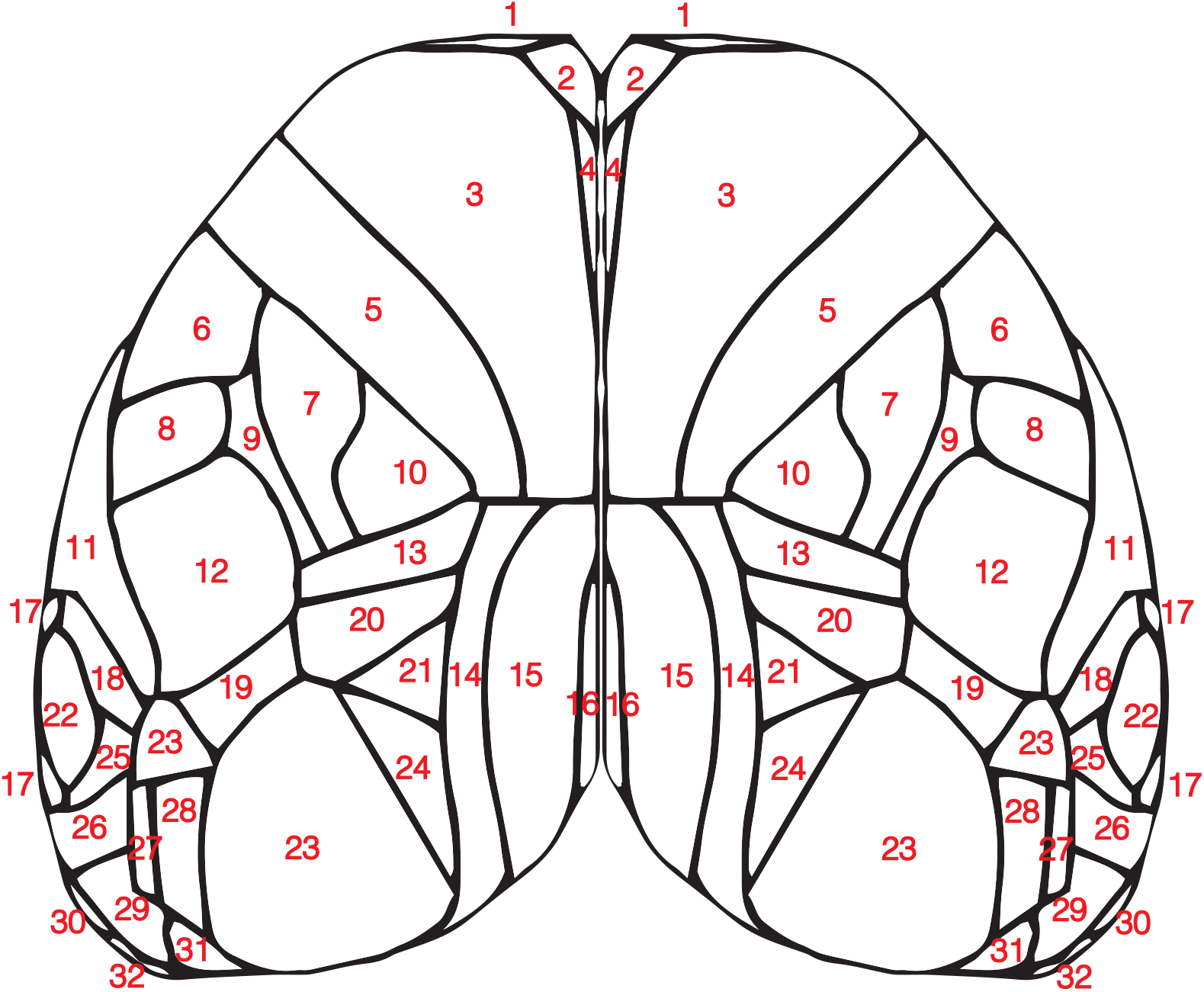
Overview of cortical areas. Cortical areas from the Allen Mouse Brain Common Coordinate Framework V3: 1. Frontal pole; 2. Prelimbic area; 3. Secondary motor area; 4. Anterior cingulate area, dorsal part; 5. Primary motor area; 6. Primary somatosensory area, mouth; 7. Primary somatosensory area, upper limb; 8. Primary somatosensory area, nose; 9. Primary somatosensory area, unassigned; 10. Primary somatosensory area, lower limb; 11. Supplemental somatosensory area; 12. Primary somatosensory area, barrel field; 13. Primary somatosensory area, trunk; 14. Retrosplenial area, lateral agranular part; 15. Retrosplenial area, dorsal part; 16. Retrosplenial area, ventral part; 17. Ventral auditory area; 18. Dorsal auditory area; 19. Rostrolateral visual area; 20. Anterior visual area; 21. Anteromedial visual area; 22. Primary auditory area; 23. Anterolateral visual area; 24. Posteromedial visual area; 25. Posterior auditory area; 26. Temporal association area; 27. Laterointermediate visual area; 28. Lateral visual area; 29. Postrhinal area; 30. Anterior cingulate area; 31. Posteriolateral visual area; 32. Entrorhinal area, medial part, dorsal zone.

**Supplementary Figure 3:**
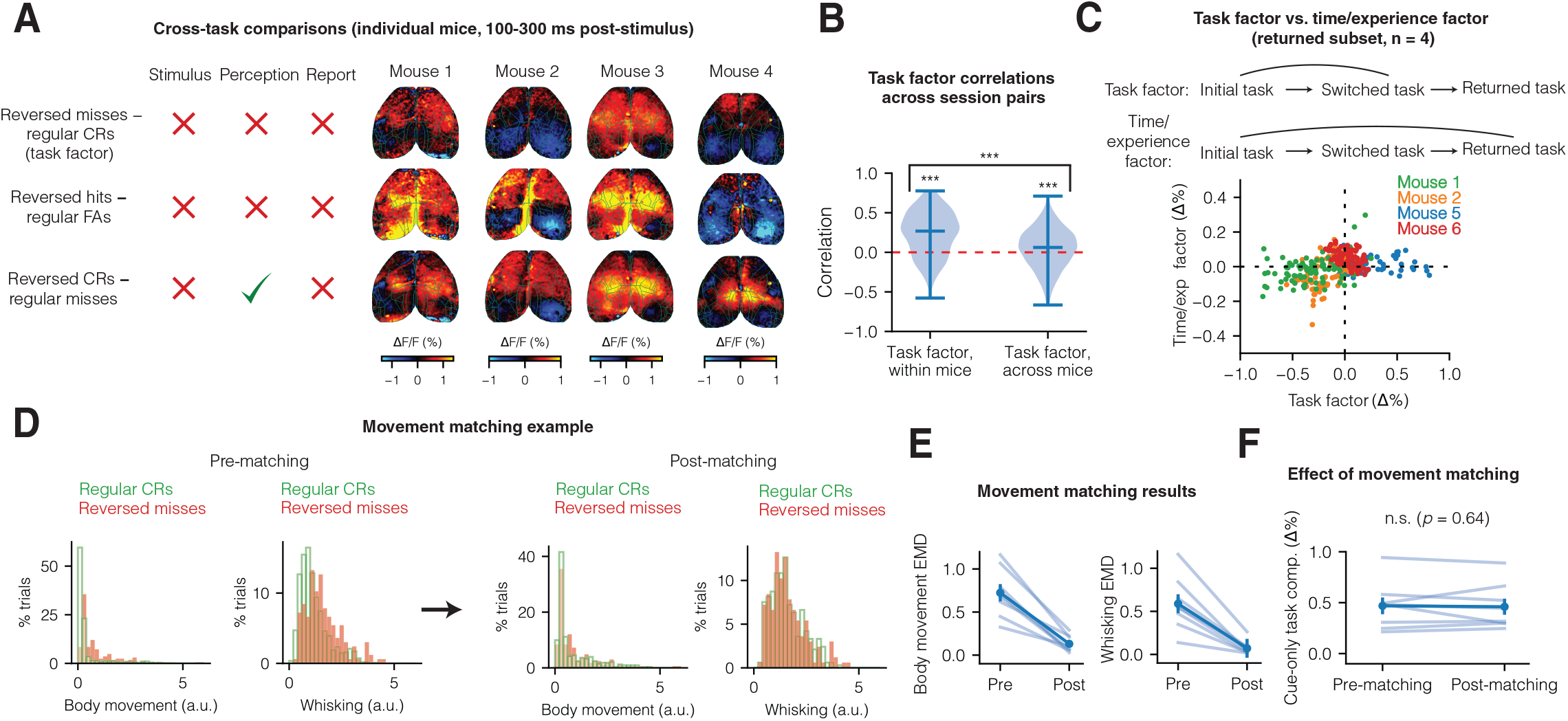
A task factor across tone-alone trials. **(A)** Individual mouse examples of the tone-alone (task factor) comparison (top row), and two others–reversed hits to regular FAs (middle row) and reversed CRs to regular misses (bottom row). **(B)** Task factor correlations across session pairs within mice vs. across mice. **(C)** Comparison of the task factor (difference in tone-alone trials between the initial and switched tasks) to the time/experience factor (difference tone-alone trials between the initial and returned task). Each dot represents a cortical region and each color a different mouse from the returned subset (n = 4 mice). **(D)** An example of the movement matching procedure for the tone-alone comparison. Pre-matching (left) and post-matching (right) histograms of mean body and whisking motion energies (MEs) during the report windows. **(E)** The overall reduction in earth mover’s distance (EMD) across the body MEs distributions (left) and whisking MEs distributions (right) due to matching (n = 8 mice). **(F)** Effect of movement matching on the mean magnitude of the task factor in each mouse. Error bars are mean ± s.e.m, *\*p* < 0.05, \*\**p* < 0.01, \*\*\**p* < 0.001.

**Supplementary Figure 4:**
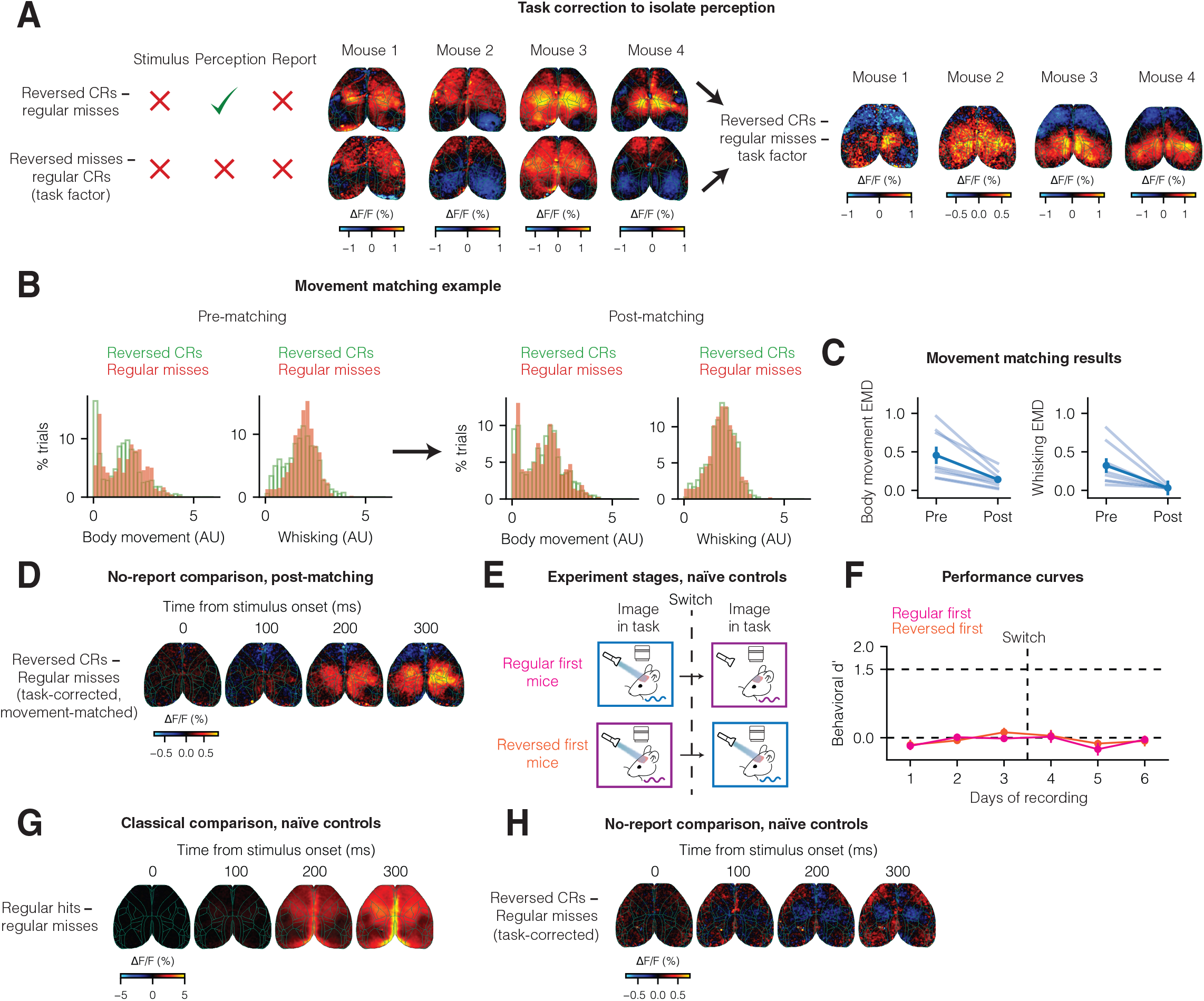
Task correction and controls. **(A)** Single mouse examples of the task correction. (Top row) The difference between reversed CRs and regular misses (no report comparison), (middle row) the difference between reversed misses and regular CRs (task factor), and (bottom row) the no report comparison subtracted by the task factor (task-corrected no report comparison). **(B)** An example of the movement matching procedure for the no-report comparison. Pre-matching (left) and post-matching (right) histograms of mean body and whisking motion energies (MEs) during the report windows. **(C)** The overall reduction in earth mover’s distance (EMD) across the body MEs distributions (left) and whisking MEs distributions (right) due to matching (n = 8 mice). **(D)** Task-corrected no-report comparison after movement matching (n = 8 mice). **(E)** Experimental stages for the n = 8 naïve control mice. **(F) (G)** Results of the classical comparison for the n = 8 naïve mice. **(H)** Results of the task-corrected no-report comparison for the n = 8 naïve mice.

